# Genomic network analysis of an environmental and livestock IncF plasmid population

**DOI:** 10.1101/2020.07.24.215889

**Authors:** William Matlock, Kevin K. Chau, Manal AbuOun, Emma Stubberfield, Leanne Barker, James Kavanagh, Hayleah Pickford, Daniel Gilson, Richard P. Smith, H. Soon Gweon, Sarah J. Hoosdally, Jeremy Swann, Robert Sebra, Mark J. Bailey, Timothy E. A. Peto, Derrick W. Crook, Muna F. Anjum, Daniel S. Read, A. Sarah Walker, Nicole Stoesser, Liam P. Shaw, on behalf of the REHAB consortium

**Author notes:** contribution considered equal (A. Sarah Walker, Nicole Stoesser, Liam P. Shaw). see acknowledgements (on behalf of the REHAB consortium). Correspondence: William Matlock and Nicole Stoesser.

## Abstract

IncF plasmids are diverse and of great clinical significance, often carrying genes conferring antimicrobial resistance (AMR) such as extended-spectrum β-lactamases, particularly in *Enterobacteriaceae*. Organising this plasmid diversity is challenging, and current knowledge is largely based on plasmids from clinical settings. Here, we present a network community analysis of a large survey of IncF plasmids from environmental (influent, effluent, and upstream/downstream waterways surrounding wastewater treatment works) and livestock settings. We use a tractable and scalable methodology to examine the relationship between plasmid metadata and network communities. This reveals how niche (sampling compartment and host genera) partition and shape plasmid diversity. We also perform pangenome-style analyses on network communities. We show that such communities define unique combinations of core genes, with limited overlap. Building plasmid phylogenies based on alignments of these core genes, we demonstrate that plasmid accessory function is closely linked to core gene content. Taken together, our results suggest that stable IncF plasmid backbone structures can persist in environmental settings while allowing dramatic variation in accessory gene content that may be linked to niche adaptation. The recent association of IncF plasmids with AMR likely reflects their suitability for rapid niche adaptation.

## Introduction

Environmental (non-clinical, non-human) populations of *Enterobacteriaceae* may act as a genetic reservoir for antimicrobial resistance (AMR). This includes livestock [1-5] and water-borne [6] resistance. Frequent horizontal gene transfer (HGT) in *Enterobacteriaceae* populations results in a large and open pangenome, enabling the wide-spread transmission of the genes conferring AMR [7-9]. This includes AMR transmission between humans and the environment and vice versa [10]. However, evidence for this transmission is often context and sequence type (ST)-specific, with broader transmission patterns less conclusive [10, 11]. IncF plasmids are a diverse group of *Enterobacteriaceae*-associated replicons which mediate the transfer of AMR genes. They have also been observed in other Proteobacteria, such as *Aeromonadaceae* and *Comamonadaceae* [12]. In particular, their involvement in the dissemination of genes encoding extended-spectrum β-lactamases (ESBLs), such as *bla-*_CTX-M-15_, is of major clinical concern [13, 14], and almost 40% of plasmid-borne carbapenemases are carried on IncF plasmids [15]. IncF plasmids are often low copy-number and conjugative [16]. Further, recent database analysis suggests IncF alleles are carried in over 50% of multireplicon plasmids [12].

Previous studies of IncF plasmids have often focussed on clinically relevant isolates, often only those encoding ESBLs [15]. Further, they have been limited to studies with smaller sample sizes. Here we analyse hundreds of IncF plasmids drawn from a survey of environmental diversity in *Enterobacteriaceae*, sampled in 2017 from a region of south-central England, UK [17]. Sampling was from livestock (cattle, pig, sheep), and from influent, effluent, and upstream/downstream waterways surrounding wastewater treatment works (collectively termed WwTWs). Potential seasonal variation was accounted for by sampling over three time-points at each site. This provided a high-quality dataset of *n*=726 plasmids for characterising natural plasmid populations.

Frequent co-integration and HGT events mean plasmid evolution cannot be described with a phylogenetic tree. Instead, networks based on sequence similarity can be used [18]. In such networks, nodes represent plasmids, and edges are weighted by a metric on the plasmid sequences. This captures both vertical and horizontal evolution at the cost of not providing a most recent common ancestor. Communities are a topological property of networks. They are defined as subsets of nodes with dense intra-connections, but sparse inter-connections [19]. In our analyses, they represented groups of similar plasmid sequences. Detecting these structures gives a coarse-grained view of the plasmid population. Previous efforts have often focussed on the relationship between network features used in plasmid classification schemes, such as replicon presence, MOB-type, or predicted mobility [20-23]. Further, studies have often focussed on curated selections from online databases [20, 22-24]. It is yet to be seen if similar community structure is present in large-scale natural populations. Additionally, it is important to develop fast and scalable methods for analysis of large and diverse WGS datasets. Here we aimed to provide a framework applicable to such studies.

## Results

### A natural population of complete plasmids with IncF replicons

We recovered *n*=726 circularised plasmids containing an IncF replicon (see Table S6) from a large dataset of high-quality *Enterobacteriaceae* genomes, obtained by hybrid assembly using both short-read (Illumina, 150bp paired end) and long-read (PacBio or Nanopore) sequencing of cultured isolates [17]. These isolates were collected over three time-points in 2017 from a region of south-central England, UK. Sampling was from 14 livestock farms (4 pig, 5 cattle, 5 sheep) and from waterways (influent, effluent and rivers) surrounding five WwTWs. Of the livestock plasmids, 120 were from pigs, 137 were from cattle and 150 were from sheep. The remaining 319 plasmids were from WwTWs.

IncF plasmids were found across all four genera in the dataset: *Citrobacter* (53 *C. freundii*), *Enterobacter* (67: 65 *E. cloacae*, 2 untyped *Enterobacter* sp.), *Escherichia* (471 *E. coli*), and *Klebsiella* (135: 61 *K. oxytoca*, 67 *K. pneumoniae*, 7 untyped *Klebsiella* sp.). Livestock plasmids mostly came from *Escherichia* (392/407), whereas WwTW plasmids had a more uniform distribution over all four genera in line with the greater diversity of genera in WwTW isolates (Fig. 1a). Our plasmids originated from *n*=558 host *Enterobacteriaceae* isolates, with a majority of chromosomes circularised (431/558).

**Figure 1.**
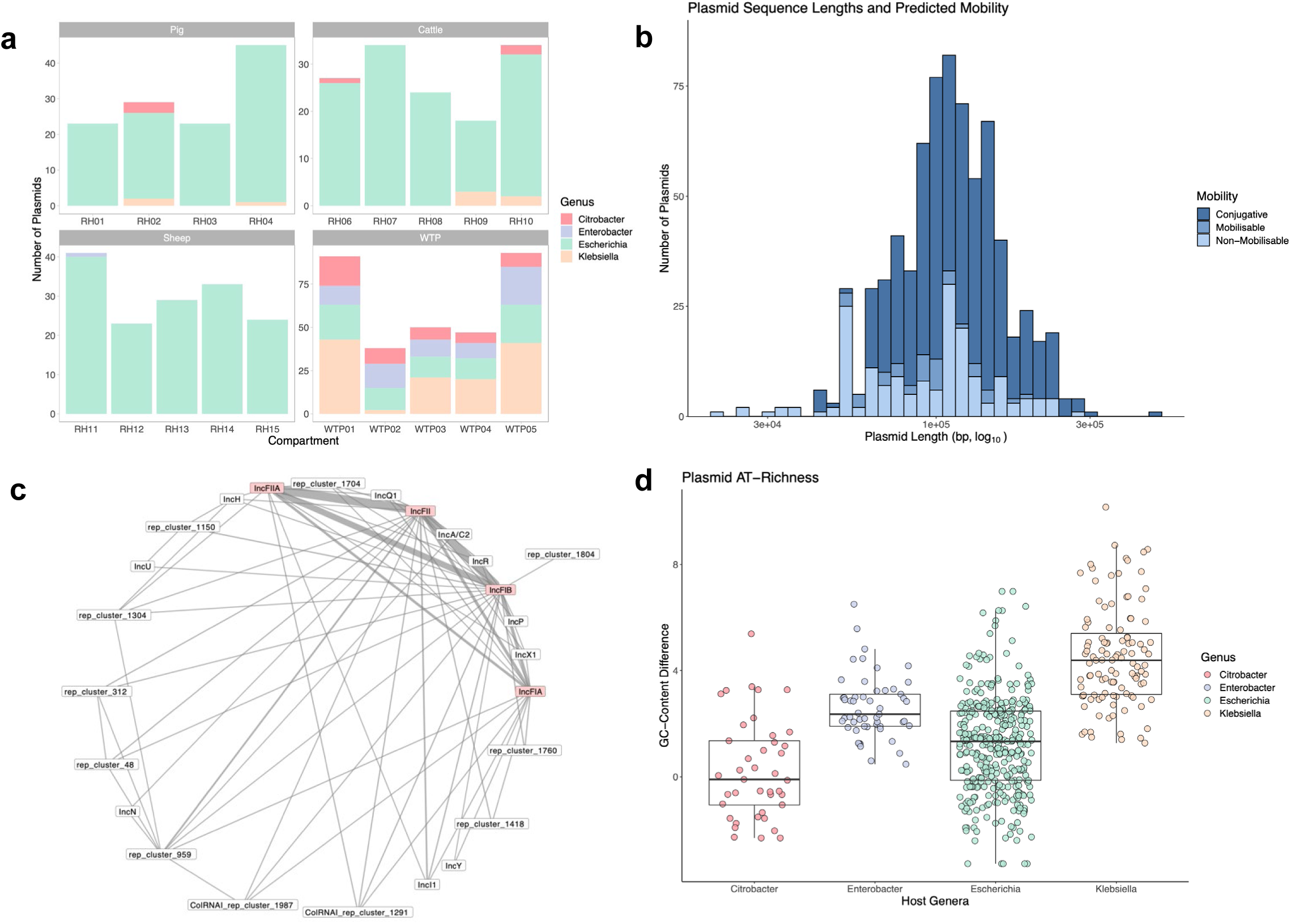
Overview of plasmid population. (a) Plasmid host genera distribution by compartment. (b) Distribution of plasmid sequence lengths with predicted mobilities. (c) Graph representing the association between replicon alleles. IncF nodes are coloured pink. Line weight is proportional to frequency of association in the sample. (d) Plasmid GC-content subtracted from host chromosome GC-content. A value greater than zero indicates the plasmid is AT-richer than the host. Only plasmids with circularised host chromosomes were used (565/726).

Plasmids ranged in length from approximately 20kb to 480kb (Fig. 1b). A majority of plasmids were predicted to be conjugative (516/726), with a smaller number predicted to be mobilisable (39/726) or non-mobilisable (171/726) (see Materials and Methods). Plasmid length was linked to mobility, with conjugative plasmids generally larger than mobilisable and non-mobilisable plasmids (Kruskal-Wallis test [*H*=36.7, *p*=1.08e-8] followed by Dunn test with Holm adjusted *p*-value [Conj—Mob: *Z*=3.45, *p*-value=1.14e-03; Conj—Non-Mob: *Z*=5.39, *p*-value=2.07e-07; Mob—Non-Mob: *Z*=-0.54, *p*-value=0.59]). We found 25 different replicons across all plasmids, including 11 in unspecified gene clusters, present in 62 different combinations or ‘replicon haplotypes’ (Table S1). 28 replicon haplotypes appeared only once in the sample. Plasmids carried between 1 and 5 replicons, with a majority carrying 2 (318/726) or 3 (209/726). Plasmid length was positively associated with number of replicons carried (one-way ANOVA test [*F*(4, 721)=5.64, *p*-value=1.8e-4] followed by Tukey’s HSD). All plasmids contained at least one IncF replicon: IncFIA (147), IncFIB (460), IncFII (574) and IncFIIA (370). Of the remaining replicons, IncI1 was most common (28), and was always found with an IncFII replicon. We observed different replicon co-occurrrence patterns (Fig. 1c), with individual IncF replicons associated with different non-IncF replicons. For instance, IncU and IncN replicons were only found with IncFIB and IncFII respectively. Overall, these co-occurrence patterns corroborate previously observed patterns of frequent IncF association with replicons such as IncI1, IncX and IncR [12].

IncF plasmids tended to be AT-rich relative to their host chromosomes. This trend has been widely reported before [25, 26]. However, we found that relative AT-richness significantly varied between host genus (one-way ANOVA test [*F*(3, 561)=111, *p*-value<2e-16] followed by Tukey’s HSD), independently of average host GC-content, with *Klebsiella* plasmids having a greater relative AT-richness than other *Enterobacteriaceae* plasmids (Fig. 1d).

### Detecting communities in plasmid k-mer networks

Plasmid sequence distances were calculated using MASH, a *k*-mer based distance estimation [27] (see Materials and Methods), and these distances used as weighted edges in a plasmid network. The output MASH edge list is presented in Table S7. Communities were detected using the Louvain algorithm, which optimises the modularity of the networks, and is a weighted community detection algorithm, meaning it also accounts for the MASH distances [19]. To effectively detect communities, we reduced the density of the network by thresholding the edges (i.e. by ‘sparsification’). Fig. 2a-b shows how the number of identified communities and percentage of plasmids covered changed as the edge (i.e. MASH distance) threshold was varied. Generally, the Louvain algorithm became more consistent in coverage as we sparsified. To ensure the communities represented potential sub-populations, we only considered those with at least 10 plasmids. The large drop in community coverage seen at threshold=0.175 (Fig. 2b) was due to the break-up of a large connected component (analogous plots for communities with at least 3 plasmid members are shown in Fig. S1a-b).

**Figure 2.**
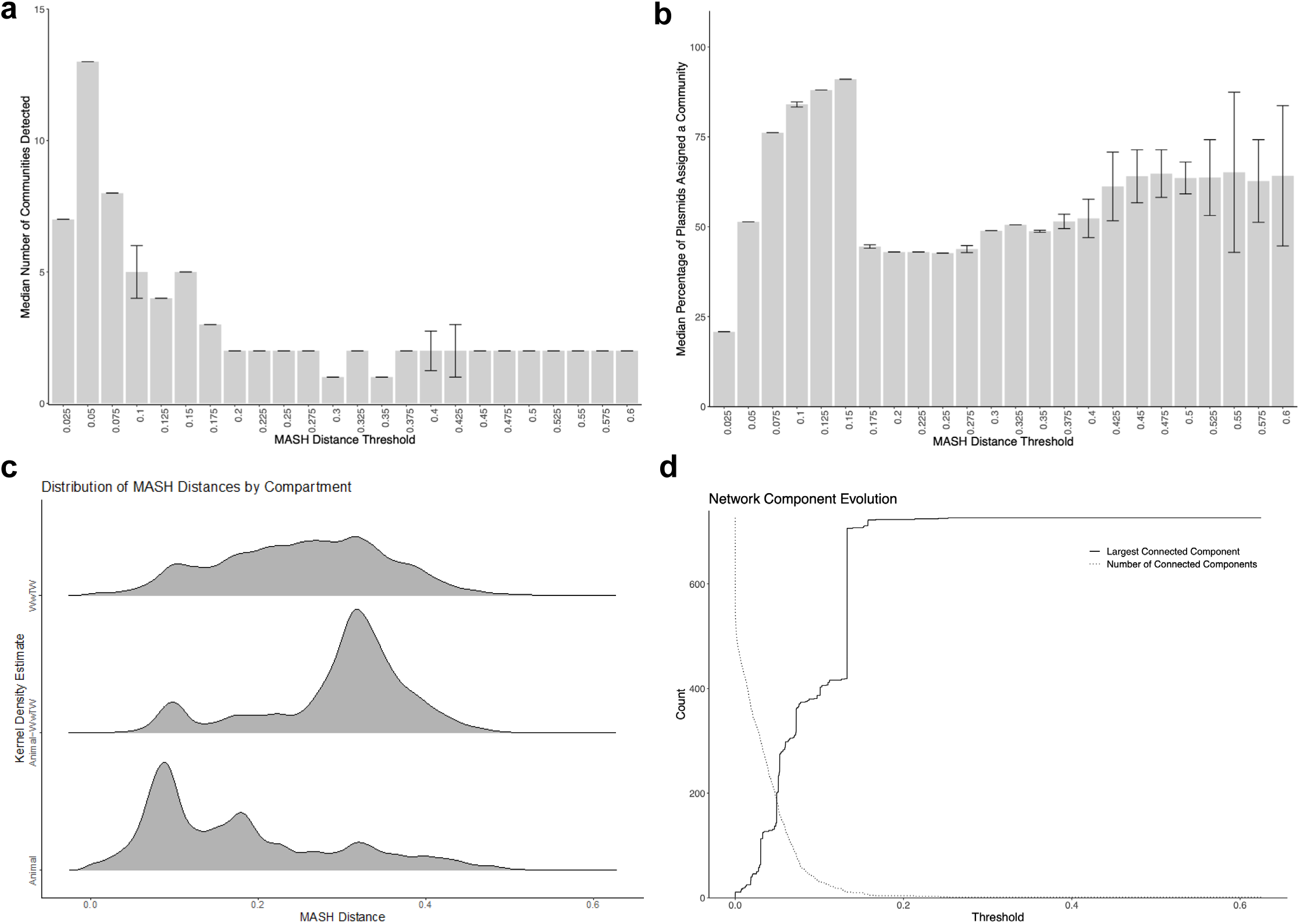
Thresholding the plasmid network. (a) Number of communities (at least 10 nodes) detected over a varying MASH threshold. Median and IQR bars shown. (b) Community coverage of the network over a varying MASH threshold. Median and IQR bars shown. (c) Gaussian kernel density estimates of MASH distances stratified by compartment. Bandwidth calculated by Silverman’s ‘rule of thumb’. (d) Evolution of the largest connected component and number of components over a varying MASH threshold.

The application of different MASH distance thresholds revealed different community structures within the network. Fig. 2c shows the kernel density estimates (KDEs) of MASH distances stratified by sampling compartment, with an overall range of [0, 0.602], highlighting that livestock plasmids (median=0.152) were generally more similar to each other than WwTW plasmids (median=0.258) and suggesting that plasmid diversity was higher in WwTW isolates. A distance threshold low enough to reveal the livestock sub-network structure could destroy the structure of the WwTW sub-network, so for this study, we selected a threshold=0.05, which revealed the structure of livestock plasmids at the expense of some WwTW structure break-up. Note that this threshold is far lower than those used in previous plasmid network analyses of global plasmid diversity [21-23]; our sample was smaller and restricted to a broad plasmid family so required more severe sparsification to reveal communities. At this level, the network’s largest connected component (LCC) had 201 nodes with 182 connected components in total (Fig. 2d). It also had the highest number of communities (13) containing at least 10 plasmids (Fig. 2a), and coverage of over 50% (Fig. 2b). There were 99 singleton plasmids, consistent with high levels of diversity in the population. A visualisation of the network at this threshold with the 13 communities coloured is presented in Fig. 3.

**Figure 3.**
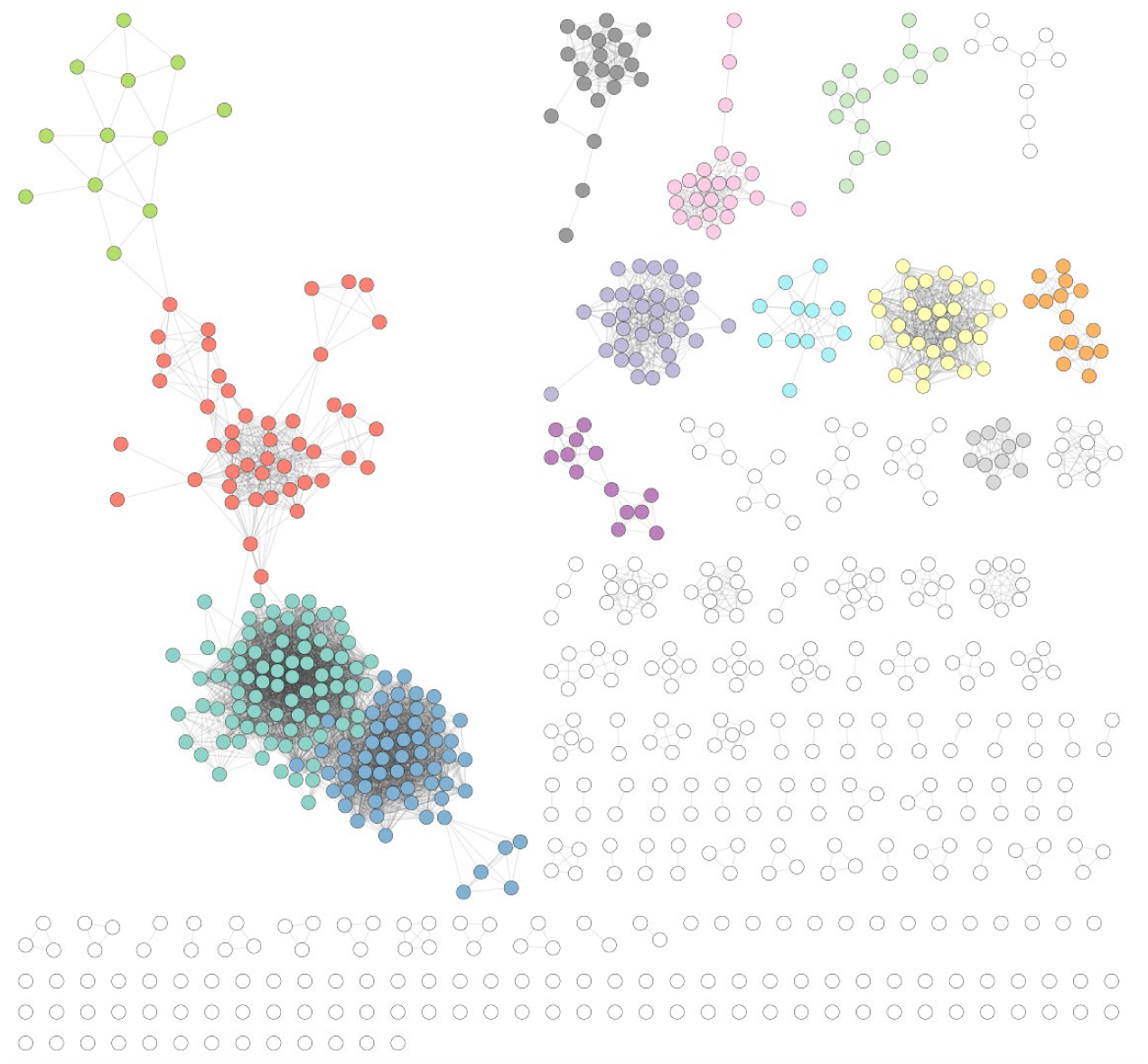
Plasmid network communities. The plasmid network at threshold=0.05. Each community with at least 10 members has unique colour. Unassigned plasmids and those in smaller communities are left white.

### Community metadata analysis

To evaluate the relationship between the node metadata labels and the network, two entropic measures were considered: homogeneity (*h*) and completeness (*c*) (both range from 0-1; see Materials and Methods). Homogeneity measures the distribution of labels given a community, with an ideal community containing a single label: a high homogeneity means that plasmids with similar sequences tend to have similar metadata labels. Conversely, completeness measures the distribution of communities given a label: a high completeness means that instances of a label tend to fall within a single community. Importantly, both homogeneity and completeness are independent of community size, the number of communities, and the number of metadata labels. This makes the approach robust to uneven sampling strategies, such as the disproportionate number of *E. coli* isolates in our sample.

Each plasmid was assigned a set of metadata labels, consisting of a sampling compartment (livestock type [pig, cattle, sheep] or WwTW-association [influent, effluent, upstream, downstream]), a host genus (*Citrobacter, Enterobacter, Escherichia* or *Klebsiella*), and a time-point (1, 2 or 3). Homogeneity (Table 1) and completeness (Table 2) were averaged over 100 runs of the Louvain algorithm. Despite the number of communities remaining consistent, some variation in the measures arose from minor changes in community boundaries.

**Table 1.** Community metadata homogeneity. Homogeneity score averages over 100 runs of the Louvain algorithm for all 13 communities.

| Table 1. Community metadata homogeneity. |  |  |  |  |  |  |  |
| --- | --- | --- | --- | --- | --- | --- | --- |
| | Mean ± sd Homogeneity | | | | | | |
| Median ± IQR<br>Communities<br>with at least 10<br>Plasmids | Livestock,<br>WwTW | Pig,<br>Cattle,<br>Sheep,<br>WwTW | 14<br>Livestock<br>Farms,<br>WwTW | Livestock,<br>5 WwTWs | Livestock,<br>Upstream/<br>Influent,<br>Downstream/<br>Effluent | Host<br>Genera | Time-point |
| 13 ± 0 | 0.715 ±<br>0.002 | 0.591 ±<br>0.008 | 0.403 ±<br>0.005 | 0.467 ±<br>0.012 | 0.550 ± 0.009 | 0.888 ±<br>0.000 | 0.050 ±<br>0.001 |

**Table 2.** Community metadata completeness. Completeness score averages over 100 runs of the Louvain algorithm for all 13 communities.

| Table 2. Community metadata completeness. |  |  |  |  |  |  |  |
| --- | --- | --- | --- | --- | --- | --- | --- |
| | Mean ± sd Completeness | | | | | | |
| Median ± IQR<br>Communities<br>with at least 10<br>Plasmids | Livestock,<br>WwTW | Pig,<br>Cattle,<br>Sheep,<br>WwTW | 14<br>Livestock<br>Farms,<br>WwTW | Livestock,<br>5 WwTWs | Livestock,<br>Upstream/<br>Influent,<br>Downstream/<br>Effluent | Host<br>Genera | Time-point |
| 13 ± 0 | 0.199 ±<br>0.002 | 0.332 ±<br>0.001 | 0.400 ±<br>0.001 | 0.238 ±<br>0.002 | 0.211 ± 0.002 | 0.307 ±<br>0.002 | 0.023 ±<br>0.000 |

Homogeneity scores showed that sampling compartment shaped plasmid similarity. At the coarsest resolution there was high homogeneity considering livestock versus WwTW (*h*=0.715; Table 1), meaning that plasmid communities were largely distinct between livestock and WwTW settings. This metadata partition is projected on the network in Fig. 4a. However, homogeneity was lower when comparing different livestock types (pig, cattle, sheep) (*h*=0.591) and even more so when comparing different farms (*h*=0.403), meaning that there was a loss of structure at these levels and plasmid communities were not well segregated by individual farm. Homogeneity was also low if plasmids were stratified by individual WwTWs (*h*=0.467). However, homogeneity increased for influent/upstream versus effluent/downstream compartments (*h*=0.550) indicating some differences in plasmids before and after WwTW treatment. Overall, plasmids from WwTWs were weakly structured by wastewater catchment.

**Figure 4.**
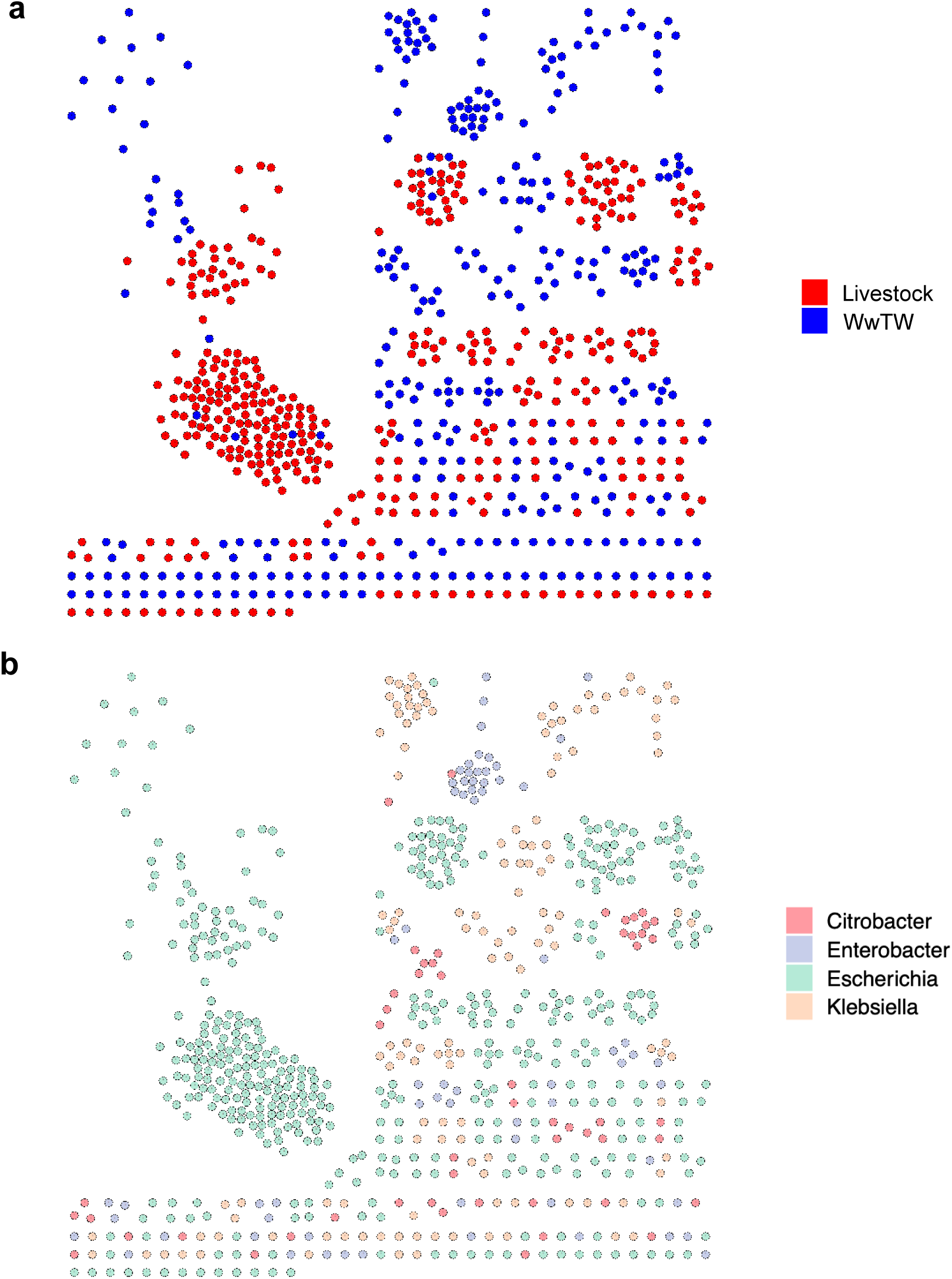
Plasmid network coloured by metadata. All nodes are coloured, not just those in our detected 13 communities of at least 10 members. (a) Partition by livestock or WwTW sampling compartment. (b) Partition by plasmid host genera.

Completeness scores highlighted higher WwTW diversity compared to lower livestock diversity. For the binary livestock or WwTW label plasmid communities scored a low completeness (Table 2; *c*=0.199), which changed little when stratified over the individual WwTWs (*c*=0.238), indicating a uniform distribution of WwTW labels over the plasmid communities and high diversity. Based on our MASH distance KDEs (Fig. 2c), we would expect livestock plasmids to have higher completeness scores than WwTW plasmids due to the lower levels of diversity; as anticipated, when stratifying the livestock metadata, completeness scores increased (*c*=0.332 and *c*=0.400). This indicated plasmids from the same farm were more likely to be found in the same community.

Host genus also played an important factor in partitioning plasmid diversity. The homogeneity scores were very high, implying a significant genetic partition by host (Table 1; *h*=0.888). This metadata partition is displayed in Fig. 4b. The lower completeness suggested a moderate level of diversity across all *Enterobacteriaceae* plasmids (Table 2; *c*=0.307). There was a very weak time-point effect found in the network (Tables 1, 2; *h*=0.050 and *c*=0.023). Under a one-tailed permutation test, all metadata label configurations except time-point had a zero *p*-value for homogeneity and completeness (Table S2; see Materials and Methods), indicating that overall, there was a significant association between niche (sampling compartment and host genus) and plasmid population structure.

### Community pangenomes

To explore the genetic structure of the communities we considered the set of all represented genes within a community, known as the pangenome (see Materials and Methods). Plasmids had a median of 35 annotated genes (range=4-112). Genes conferring AMR were found in 17% (122/726) of plasmids; this included 33 plasmids carrying ESBLs (9 pig, 8 cattle and 16 WwTw), with 4 carrying *bla*_CTX-M-15_ (all WwTW). IncF plasmids in isolates cultured from pigs were disproportionately associated with AMR genes (45/109 [41%] AMR plasmids).

Core genes with well-conserved synteny comprise the plasmid ‘backbone’ [18], which often controls essential replication and mobility functions. Genes with accessory function, such as AMR genes, are inserted into the backbone. For the 13 IncF plasmid communities identified in this study using the 0.05 threshold above (see Fig. 3), we found a median of 13 core genes (range=0-88) (Table 3). Each community possessed a unique combination of core genes, and pairs of communities shared a median of 0 core genes between them (range=0-21) (Table S3). The communities had a median of 463 accessory genes (range=151-790), sharing a median of 312 accessory genes (range=99-570) (Table S4). Pairs of communities sharing a higher number of genes tended to have a higher sum of individual genes (*r*=0.820, *t*=12.505, *p*-value<2.2e-16), indicating overlap between larger pangenomes. Within a plasmid community, a lower mean MASH value indicates greater overall sequence similarity; as anticipated therefore, we found a lower mean MASH distance was associated with more core genes (*r*=-0.615, *t*=-2.586, *p*-value=0.025) and a lower total number of genes in the pangenome (*r*=0.654, *t*=2.865, *p*-value=0.015).

**Table 3.** Community pangenome results. Characteristics of each of the 13 communities, including number of nodes, edges and MASH mean (mean weight of all edges), and gene counts at each level of the pangenome: core genes, soft core genes, shell genes and cloud genes are those found in [100, 99], (99, 95], (95, 15], and (15, 0] percent of plasmids respectively.

| **Table 3. Community pangenome results.** |  |  |  |  |  |  |  |  |
| --- | --- | --- | --- | --- | --- | --- | --- | --- |
| **Community** | **Nodes** | **Edges** | **MASH mean** | **Core Genes** | **Soft Core Genes** | **Shell Genes** | **Cloud Genes** | **Total Genes** |
| 1 | 61 | 1252 | 0.0284 | 3 | 6 | 172 | 263 | 444 |
| 2 | 82 | 1817 | 0.0314 | 4 | 18 | 138 | 364 | 524 |
| 3 | 46 | 325 | 0.0355 | 35 | 8 | 86 | 369 | 498 |
| 4 | 12 | 21 | 0.0382 | 2 | 0 | 290 | 129 | 421 |
| 5 | 14 | 23 | 0.0383 | 2 | 0 | 225 | 260 | 487 |
| 6 | 21 | 111 | 0.0366 | 13 | 6 | 354 | 430 | 803 |
| 7 | 34 | 263 | 0.0344 | 2 | 1 | 278 | 359 | 640 |
| 8 | 23 | 135 | 0.0222 | 27 | 1 | 142 | 362 | 532 |
| 9 | 12 | 34 | 0.0344 | 18 | 0 | 364 | 324 | 706 |
| 10 | 13 | 37 | 0.0233 | 0 | 0 | 309 | 38 | 347 |
| 11 | 15 | 55 | 0.0194 | 62 | 0 | 116 | 35 | 213 |
| 12 | 30 | 391 | 0.0242 | 68 | 3 | 126 | 187 | 384 |
| 13 | 12 | 45 | 0.0223 | 88 | 0 | 195 | 48 | 331 |

For an example community of 30 IncF plasmids from isolates from sheep farms, we produced a neighbour-joining phylogeny based on 64/384 core genes (Fig. 5). The tree accounts for homologous recombination, with events detected in 11/30 leaf nodes and 21 internal nodes, consistent with a high number of exchange events affecting this plasmid community. The median tract length was 156bp (range=2bp-2249bp). Annotation of the phylogeny with the 316 accessory genes for this community revealed that accessory gene presence aligned almost identically with the core gene phylogeny, suggesting that the evolution of the plasmid backbone is highly linked to accessory function. All host genera for this plasmid community were diverse *E. coli*, with 13 known STs present, consistent with widespread horizontal transfer of the plasmids from this community. Within this community, no plasmids carried AMR genes. Core genome phylogenies for other plasmid communities also showed a strong link between accessory gene presence and backbone contents (Figs. S2-S12).

**Figure 5.**
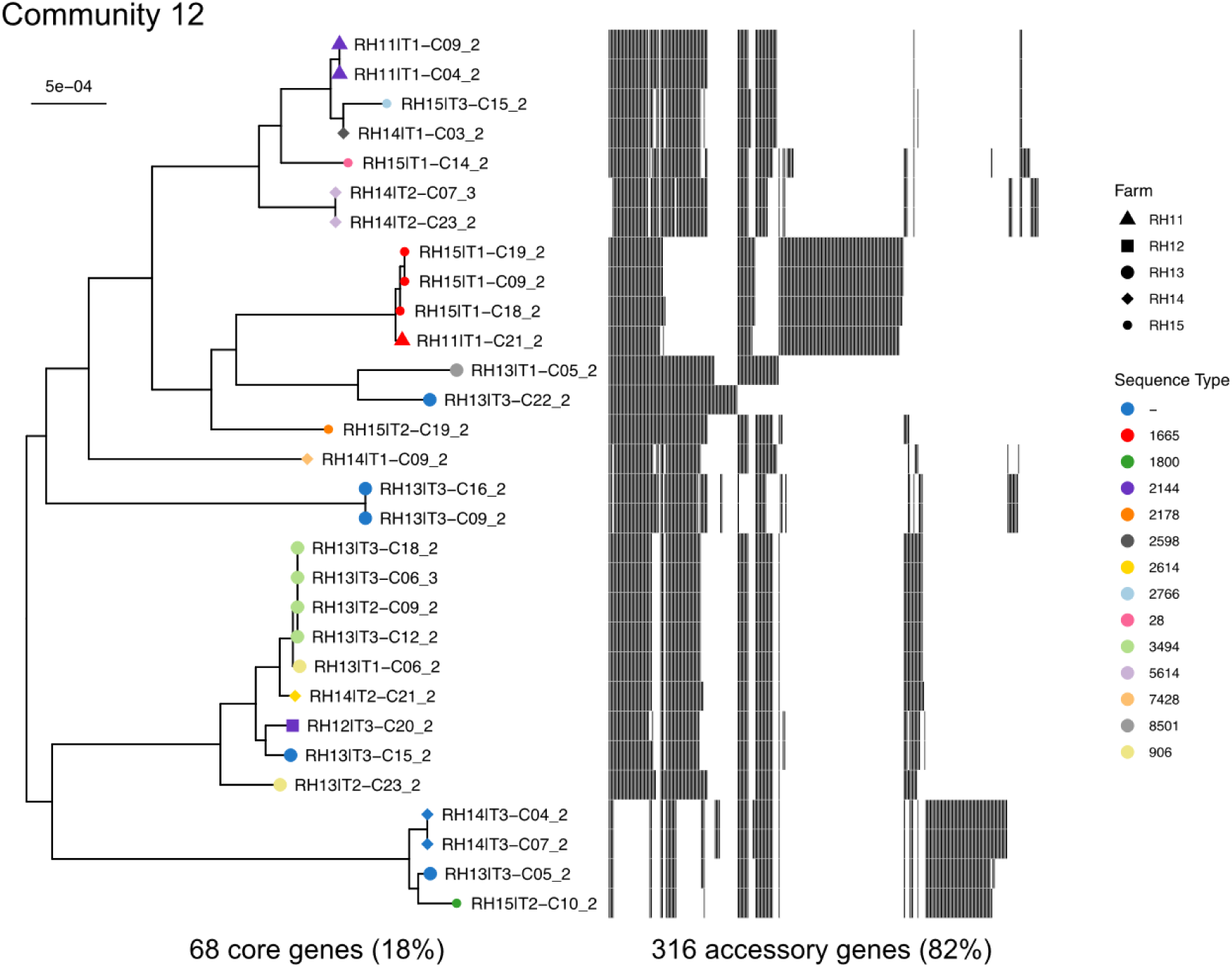
Community core gene phylogeny. A neighbour-joining tree based on alignments of the 68 core genes. A heatmap of the 316 accessory genes is also shown. Node colour represents host sequence type and node shape represents farm. Unknown STs are labelled by ‘-’. Branch lengths have been corrected for homologous recombination.

## Discussion

We have analysed plasmid communities using distance-free genomic networks to explore diversity within a large, natural population of IncF plasmids from four *Enterobacteriaceae* genera (*Citrobacter, Enterobacter, Escherichia* and *Klebsiella*). These IncF plasmids contained a diversity of replicons (plasmids contained 21 other replicons, forming 62 unique combinations) and we resolved plasmids into communities (12 communities of ≥10 plasmids). We found that 15% of IncF plasmids contained at least one AMR gene, and 5% carried an ESBL. This underlines that non-clinical plasmid populations can also carry AMR genes, and that WwTW environment and livestock niches are part of an AMR network for *Enterobacteriacae* [2, 10].

Our network analysis revealed IncF plasmids were well partitioned by sampling compartment, with distinct communities isolated to WwTWs or livestock; however, there were also clear instances of sharing events between, for example, specific farm locations. There was also moderate partitioning by specific livestock species: pig, cattle and sheep. Additionally, there was a difference in plasmids before and after WwTW treatment. Sampling compartment also influenced diversity, with a higher diversity in WwTW-associated plasmids than livestock plasmids. This is probably because both river and wastewater catchments integrate a large number of human, livestock (farmed and wild) and environmental sources. Further, they also experience higher rates of inflow and outflow than farm-specific environments. The analysis also revealed a significant partition by host genera. Despite IncF plasmids ranging over all *Enterobacteriaceae* genera, it suggested some genus-specific adaptations. Notably, the extent of plasmid-host AT-richness relative to the host chromosome varied depending on the genus. It remains to be seen how such observed differences relate to plasmid function. However, this may be related to the livestock-WwTW partition, since our livestock plasmids were predominantly hosted by *E. coli*. We did not detect an effect of sampling time-point. This is may be because our time-points were too close and sample size too small to capture any significant evolution, or it may indicate that time of year is not a strong factor in determining community structure. It would be interesting to see how plasmids from clinical samples relate to those from our samples within the network, especially if pre-WwTW plasmids are considered as a proxy for human gut microbiomes.

Pangenome analysis of the inferred plasmid communities revealed that core gene content was mostly unique to communities. Further, they were strongly related to accessory function. Taken with the above results, we propose that sampling compartment and host greatly influence the function of plasmids. This includes AMR presence, with pigs, and hence *Escherichia*, carrying a disproportionate burden in our sample. The pangenomes for communities varied greatly in the number of core genes, with one community having zero. This may be because the threshold was not severe enough to segregate this particular community into uniquely similar groups. It also may result from how Panaroo (see Materials and Methods) corrects annotation errors, splitting gene clusters too readily. Generally, more genetically similar communities possessed a greater number of core genes and smaller pangenome. Our results for IncF plasmid communities are in line with a recent study of the wider prokaryotic plasmidome which concluded that clusters of plasmids contain common genomic backbones [23].

Our study has several limitations. One important limitation, which applies more widely to network approaches which cluster or partition diversity, is that thresholding of the network is highly subjective and dataset dependent. Trade-offs are required to reveal the intermediate structures of the network whilst maintaining good community detection performance. We determined a threshold by considering MASH distance distributions and component evolution alongside Louvain output diagnostics. When diversity varies greatly between sampling compartments, a single threshold is unlikely to be globally optimal. In these cases, it is probably best to focus on sub-populations of interest. Despite only considering several hundred nodes here, our methodology is scalable to far larger studies. Originally, the Louvain algorithm had runtime complexity *O*(*e*), where *e* is the number of edges in the network. This has since been improved to *O*(*v*log*k*), where *v* is the number of nodes and *k* is the average node degree [28]. Further, recent efforts have parallelised the Louvain algorithm to networks with billions of edges, though this approach was not necessary here [29]. Finally, our dataset is limited to the four *Enterobacteriaceae* genera under study and conclusions may not reflect the wider diversity of IncF plasmids beyond these genera.

In conclusion, our study adds to the growing literature on distance-free networks to characterise and partition plasmid diversity, introducing a scalable framework to quantify the relationship between network structure and plasmid metadata by identifying network communities. Overall, our approach represented a high-resolution strategy for summarising similarities and differences within plasmid populations, leveraging the advantages of having complete plasmid sequences and analysing these in the context of associated metadata. For IncF plasmids we were able to show the distinct, local effects of sampling compartment on plasmid structure and population, but also identify evidence for sharing of plasmids between bacterial lineages, farms and WwTW-associated contexts, with relevance for the “One Health”-associated study of mobile genetic elements and AMR genes. As long-read sequencing costs fall, and increasingly large numbers of plasmids can be characterised, future work applying this method will contribute to better understanding plasmid populations, estimating transfer rates of important AMR genes and MGEs between potential reservoirs, and identifying hotspots of selection/transfer that might be amenable to intervention.

## Materials and Methods

Plasmids and corresponding host isolates were sampled and sequenced on behalf of the REHAB project in 2017, which aimed to characterise the non-clinical, non-human *Enterobacteriaceae* microbiome in south-central England, with a focus on better understanding antimicrobial resistance (AMR) spread. Specifically, livestock (pig farms, cattle farms and sheep farms) and wastewater treatment work environments (WwTWs; influent, effluent, upstream and downstream waterways) were sampled. To account for seasonal variation, sampling occurred at three discrete time-points (TPs): January-April 2017 (TP1), June-July 2017 (TP2), October-November 2017 (TP3). All the plasmids presented have at least one IncF replicon (classified by with MOB-typer, see below). In total, we present *n*=726 plasmids originated from *n*=558 isolates. This comprises a subset of the entire REHAB dataset, which overall contains *n*=2,293 circularised plasmids recovered from *n*=828 isolates. This dataset is described in more detail [17].

### Livestock

Four pig farms (RH01-04), five cattle (RH06-10) and five sheep farms (RH11-15) were selected for sampling over all three TPs. All participating farmers provided written consent for participation. Specific details on farm recruitment and sampling procedure can be found in [17] and Anjum et al. (paper in preparation).

### Wastewater treatment works (WwTWs) environment

Five WwTWs (WTP01-05) were selected based on a number of criteria, including; geographic location within the region, wastewater treatment configuration, wastewater population equivalent (PE) served, consented flow, and the accessibility of the effluent receiving river for sampling both upstream and downstream. The chosen WwTWs and their details are shown in Table S5. Sampling took place over all three TPs. Specific details are provided in [17].

### DNA sequencing

The isolates were selected for sequencing to represent diversity within the four major genera (*Citrobacter, Enterobacter, Escherichia* and *Klebsiella*) in each niche, including the use of third-generation cephalosporin resistance to identify a subset of multi-drug resistant isolates within each genus. Sequencing involved either PacBio SMRT (*n=*293) or Oxford Nanopore Technologies (ONT) (*n*=268) methodologies. Specific details are provided in [17].

### Genome assembly, assignment and typing

We used the hybrid assembly and sequencing methods described in our pilot study [30] to produce high-quality *Enterobacteriaceae* genomes from short and long reads. We assigned species and sequence type (ST) from assembled genomes using mlst (version 2.16.43) [31]. Further details on validation are provided in [17].

### Plasmid assembly

We used the hybrid assembly and sequencing methods described in a pilot study [30] to produce high-quality *Enterobacteriaceae* genomes from short and long reads. In short, we used Unicycler (version 0.4.7) [32] with ‘normal’ mode, --min_component_size 500, --min_dead_end_size 500, and otherwise default parameters. From these, we selected *n*=726 plasmids which contained an IncF replicon after classification with MOB-typer (see below). We searched all plasmids against PLSDB (version 2020-03-04) [33] which contains 20,668 complete published plasmids, using mash screen [34] and keeping the top hit. All plasmids had a match.

### Mobility typing

We used MOB-typer from MOB-suite (version 2.0.0) [35]. We clustered plasmids using MOB-cluster and assigned replicon types with MOB-typer, both part of the MOB-suite. MOB-cluster uses single linkage clustering with a cutoff of a mash distance of 0.05 (corresponding to 95% ANI). A recent large-scale study [12] showed MOB-typer to have a higher correct classification rate than the widely used PlasmidFinder [36].

### Plasmid distance estimation

Distances between the complete plasmid sequences was calculated using MASH (version 2.2) [27]. MASH reduces sequences to a fixed-length MinHash sketch, which is used to estimate the Jaccard index. This measures extent of *k*-mer sharing between plasmids. The representative sketch is far shorter than the original sequence, making distance calculations efficient over large datasets. A *k*-mer length of 13 and a sketch size of 5000 was used. All other settings were default. Using MASH considerably reduces distance computation time from exact *k*-mer profile methods, whilst maintaining good performance.

### Louvain community detection

The Louvain algorithm detects communities by optimising the modularity by iterative expectation-maximisation (EM) [19]. This aims to maximise the density of edges within communities against edges between communities. The algorithm was implemented using the python-louvain (version 0.14) Python module.

### Community metadata analysis

Homogeneity (*h*) and completeness (*c*) are dual conditional entropy-based measures [37]. They are independent of clustering algorithm, dataset size, number of label-types, number of communities and community sizes. This means they are appropriate for uneven metadata distributions. A community partition satisfies homogeneity (*h* = 1) if all members have the same metadata label-type. Suppose we have network with *N* nodes, partitioned by a set of metadata labels, *M* = {*m*_*i*_ |*i* = 1, *…, n*}, and a set of communities, *C* = {*c*_*j*_ |*j* = 1, *…, m*}. Let *A* = {*a*_*ij*_} represent the *ij*^*th*^ entry in the contingency table of partitions. Hence, *a*_*ij*_ counts the number of nodes with label *m*_*i*_ in community *c*_*j*_. We then say

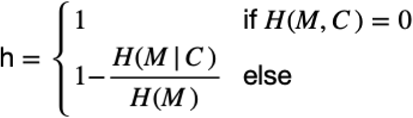

where

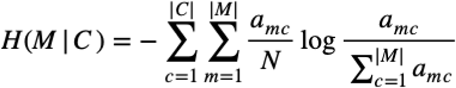

and

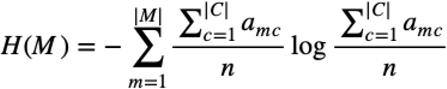

are the conditional entropy of the metadata given the communities and the entropy of the communities, respectively *H*(*M*|*C*) = 0 when the community partition coincides with the metadata partition, and no new information is added. A community partition satisfies completeness (*c* =1) if all instances of a metadata label-type are assigned the same community. Completeness is defined dually by

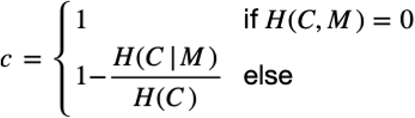

The measures were calculated using the scikit-learn (version 0.22.2) Python module [38].

### Permutation test

We first calculated a Louvain partition for the network and selected all nodes in communities with at least 10 members. Homogeneity and completeness score medians were used from Table 1 and Table 2. The partition labels were then randomly permuted 1,000 times. For each permutation, the homogeneity and completeness scores were calculated. These were then used to calculate a right-tailed *p*-value. The results are shown in Table S2.

### Plasmid annotation and pangenome analysis

Plasmids were annotated using Prokka (version 1.14.6) [39]. Pangenome analysis used Panaroo (version 1.2.2) [40]. Core genes, soft core genes, shell genes and cloud genes are those found in [100, 99], (99, 95], (95, 15], and (15, 0] percent of sequences respectively. Within the pangenome, core genes are typically defined as those shared by ≥99% of constituent plasmids. However, since no plasmid community in this study had >100 members, core genes were strictly shared by 100%. AMR annotations used Abricate (version 0.9.8) [41] with with the NCBI AMRFinder Plus database [42] with a threshold of 90% sequence identity and 90% coverage.

### Community phylogeny

Alignment of core genes used Clustal Omega (version 1.2.4) [43], and ClonalFrameML (version 1.2) [44] was used to adjust for homologous recombination. We used ggtree (version 3.11) [45] to visualise the phylogeny.

### Data visualization

All figures were made in using the R package ggplot2 (version 3.3.0) [46], except for the network figures (1c, 3 and 4a-b), which were made using Cytoscape (version 3.8.0) [47]. Cytoscape was also used to calculate some network descriptive statistics.

### Code and data availability

Plasmid sequence data, metadata (Table S6) and MASH edge list (Table S7) output are available in a figshare collection (https://doi.org/10.6084/m9.figshare.c.5066684.v1). Further details on computing methods can be found in the GitHub repository for the paper (https://github.com/wtmatlock/plasmid-network-analysis). This includes scripts for calculating the LCC and NCCs, Louvain performance diagnostics, and running the permutation test. Other data can be found in [17].

## Acknowledgements

The REHAB consortium is represented by the following: AbuOun M, Anjum MF, Bailey MJ, Brett H, Bowes M, Chau KK, Crook DW, de Maio N, Duggett N, Wilson DJ, Gilson D, Gweon HS, Hubbard A, Hoosdally SJ, Matlock W, Kavanagh J, Jones H, Peto TEA, Read DS, Sebra R, Shaw LP, Sheppard AE, Smith R, Stubberfield E, Stoesser N, Swann J, Walker AS, Woodford N. Also, thanks to Fowler P for his comments on the draft.

## Author contributions

Author contributions under the CRediT system were as follows:

Conceptualisation: WM, NS, MA, DS, MJB, DWC, LPS, ASW

Methodology: WM, LPS

Software: WM

Validation: WM, KKC, LB, HP, LPS

Formal analysis: WM

Investigation: KKC, MA, ES, JK, HP, LB, RS, DSR, HSG, NS, RS

Resources: MA, MFA, HSG, DSR, RS, JS, NS, TEAP, MJB, ASW, RS

Data curation: WM, LPS, DSR, MA, NS, ES, DG

Writing – original draft: WM

Writing – review and editing: All authors

Visualisation: WM

Supervision: LPS, NS, ASW, DWC

Project administration: NS, DSR, SH, MFA

Funding acquisition: NS, DWC, MJB, DSR, MFA, ASW, TEAP

## Competing interest declarations

The authors declare no competing interests.

## Funding

This work was funded by the Antimicrobial Resistance Cross-council Initiative supported by the seven research councils [grant NE/N019989/1]. Crook, George, Peto, Sheppard, Stoesser, and Walker are supported by the National Institute for Health Research Health Protection Research Unit (NIHR HPRU) in Healthcare Associated Infections and Antimicrobial Resistance at the University of Oxford in partnership with Public Health England (PHE) [grant HPRU-2012–10041 and NIHR200915]. Walker, Crook, and Peto are also supported by the NIHR Oxford Biomedical Research Centre. Walker is an NIHR Senior Investigator. The computational aspects of this research were funded from the NIHR Oxford BRC with additional support from a Wellcome Trust Core Award Grant [grant 203141/Z/16/Z]. The views expressed are those of the authors and not necessarily those of the NHS, the NIHR, the Department of Health or Public Health England. Matlock is supported by a scholarship from the Medical Research Foundation National PhD Training Programme in Antimicrobial Resistance Research (MRF-145-0004-TPG-AVISO).

## Notes

### Competing Interest Statement

The authors have declared no competing interest.

