## Supplementary Materials for "Genomic network analysis of an environmental and livestock IncF plasmid population"

**Supplementary figures:**
Figure S1: Louvain performance for communities with at least 3 members.
Figures S2-S-12: Plasmid core-gene phylogenies for communities 1, 2, 3, 4, 5, 6, 7, 8, 9, 11, and 13

**Supplementary tables:**
Table S1: 'Replicon haplotype' counts for the *n*=726 IncF plasmids
Table S2: *p*-values for the permutation test on homogeneity and completeness scores
Table S3: Matrix counting the number of shared core genes between communities 1-13
Table S4: Matrix counting the number of shared accessory (non-core) genes between communities 1-13
Table S5: Metadata for the 5 WwTW sampling locations

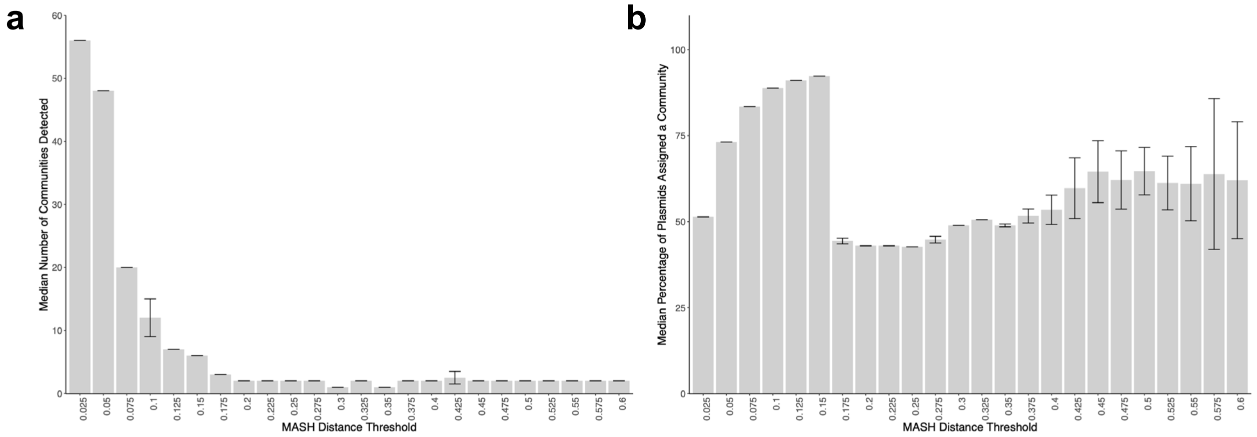


**Figure S1. Louvain performance for communities with at least 3 members.** (a) Number of communities detected over a varying MASH threshold. (b) Community coverage of the network over a varying MASH threshold.

### Community 1

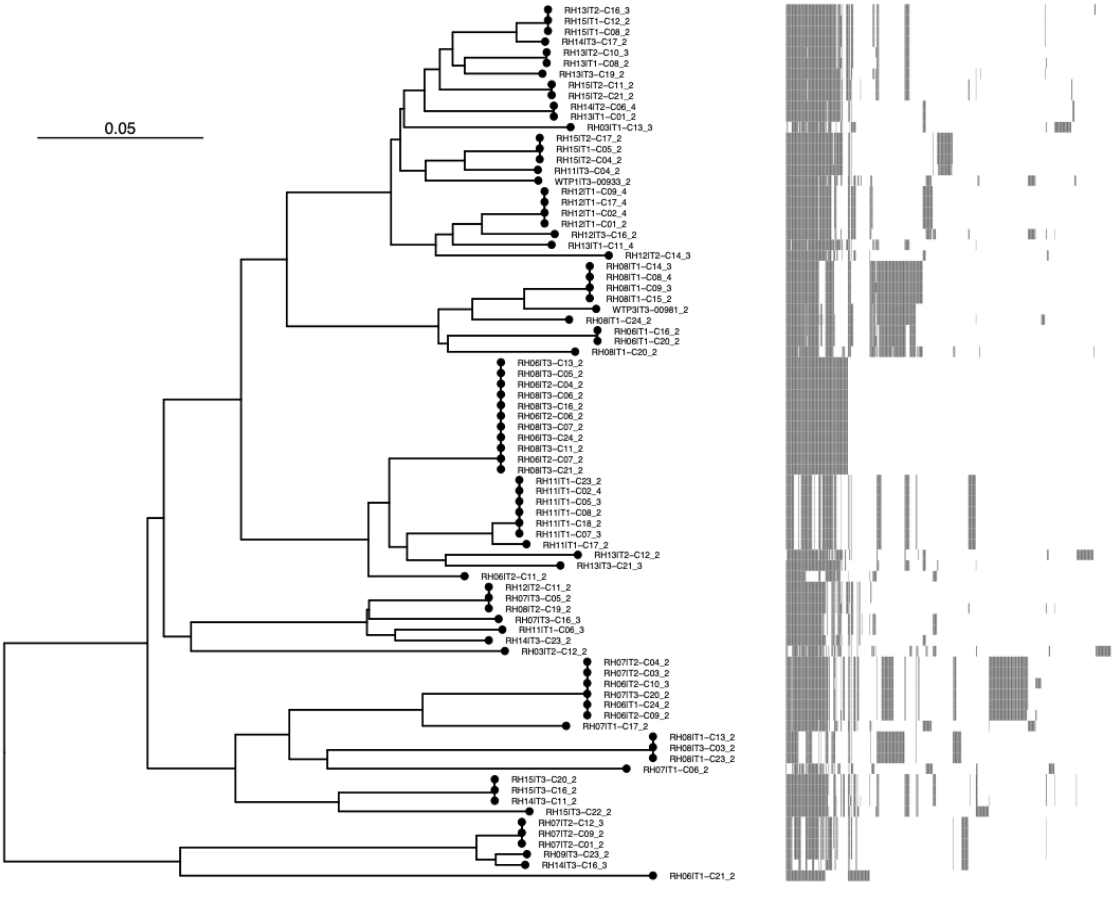


**Figure S2. Plasmid core-gene phylogeny for community 1.**

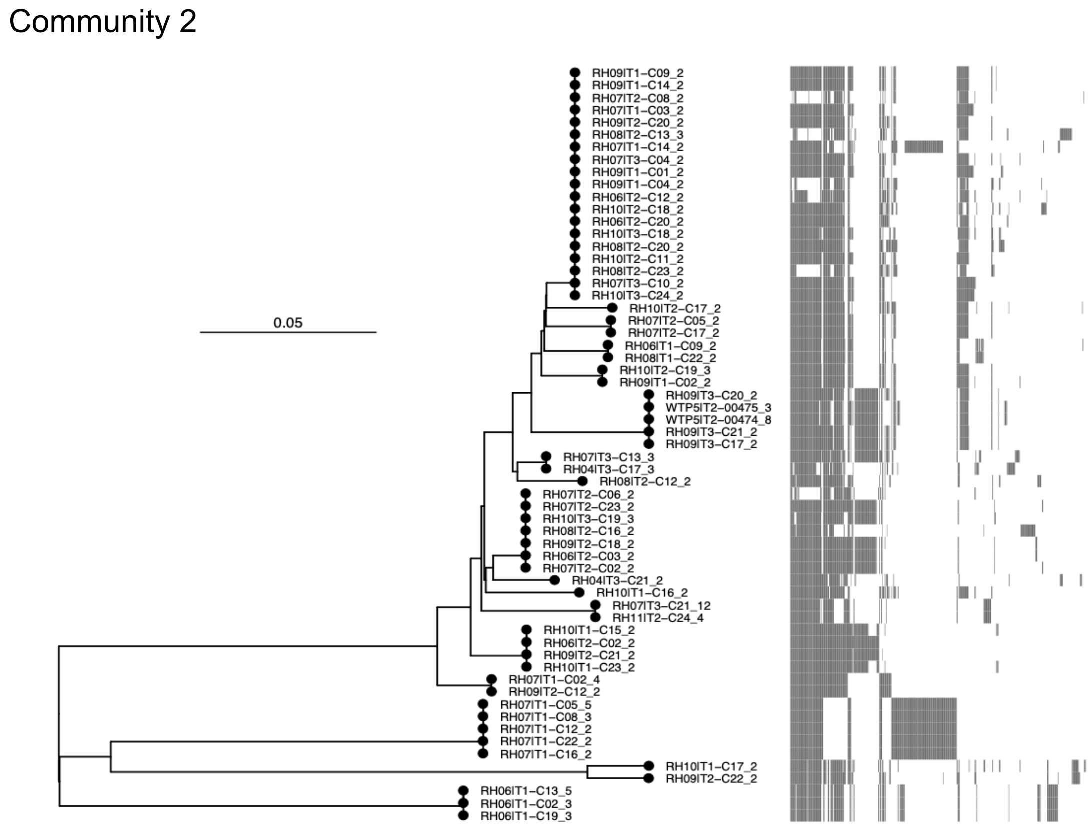


**Figure S3. Plasmid core-gene phylogeny for community 2.**

Community 3

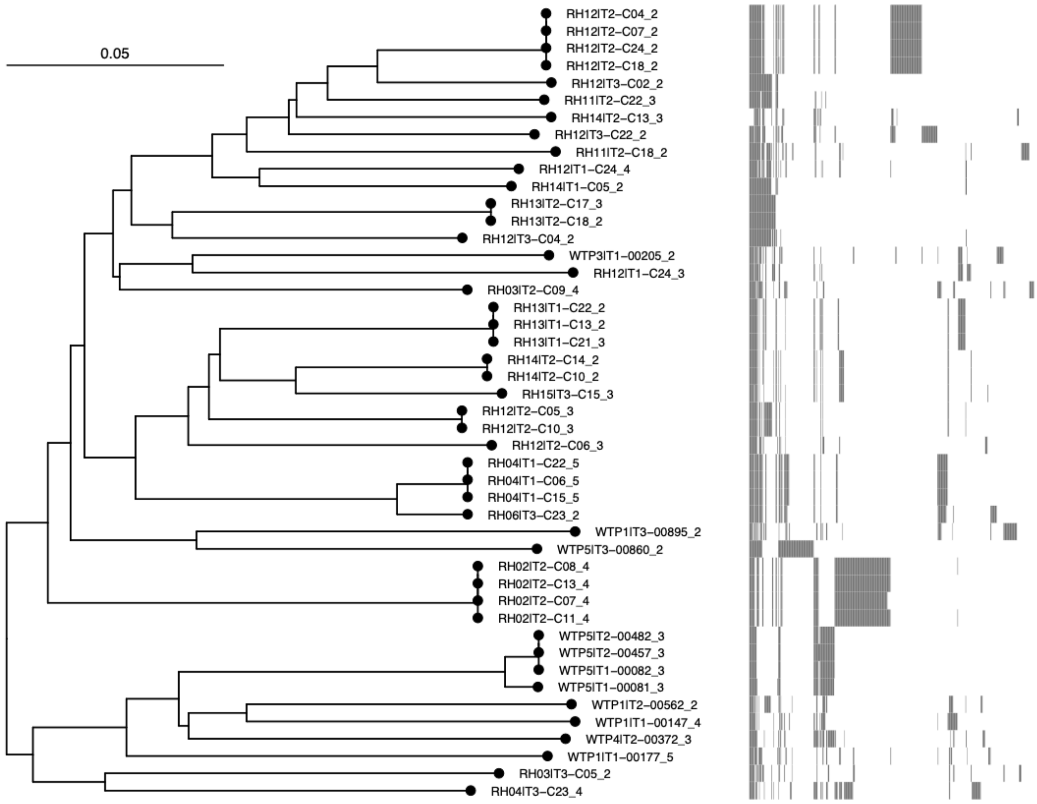


**Figure S4. Plasmid core-gene phylogeny for community 3.**

Community 4

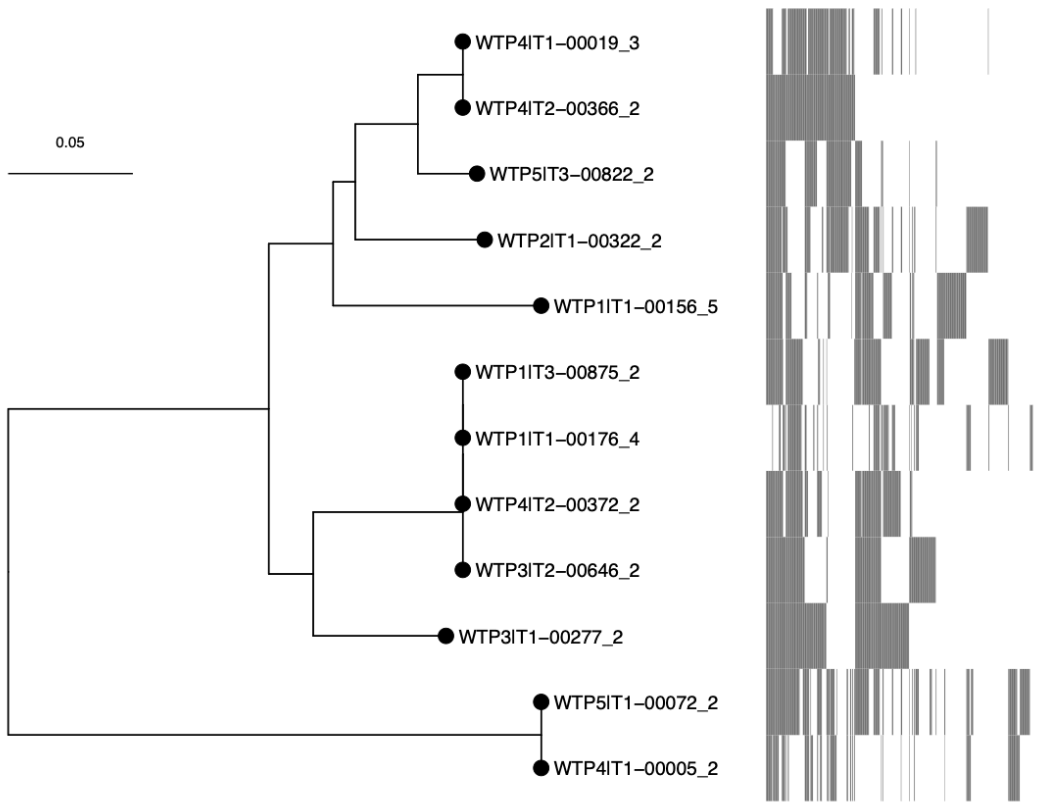


**Figure S5. Plasmid core-gene phylogeny for community 4.**

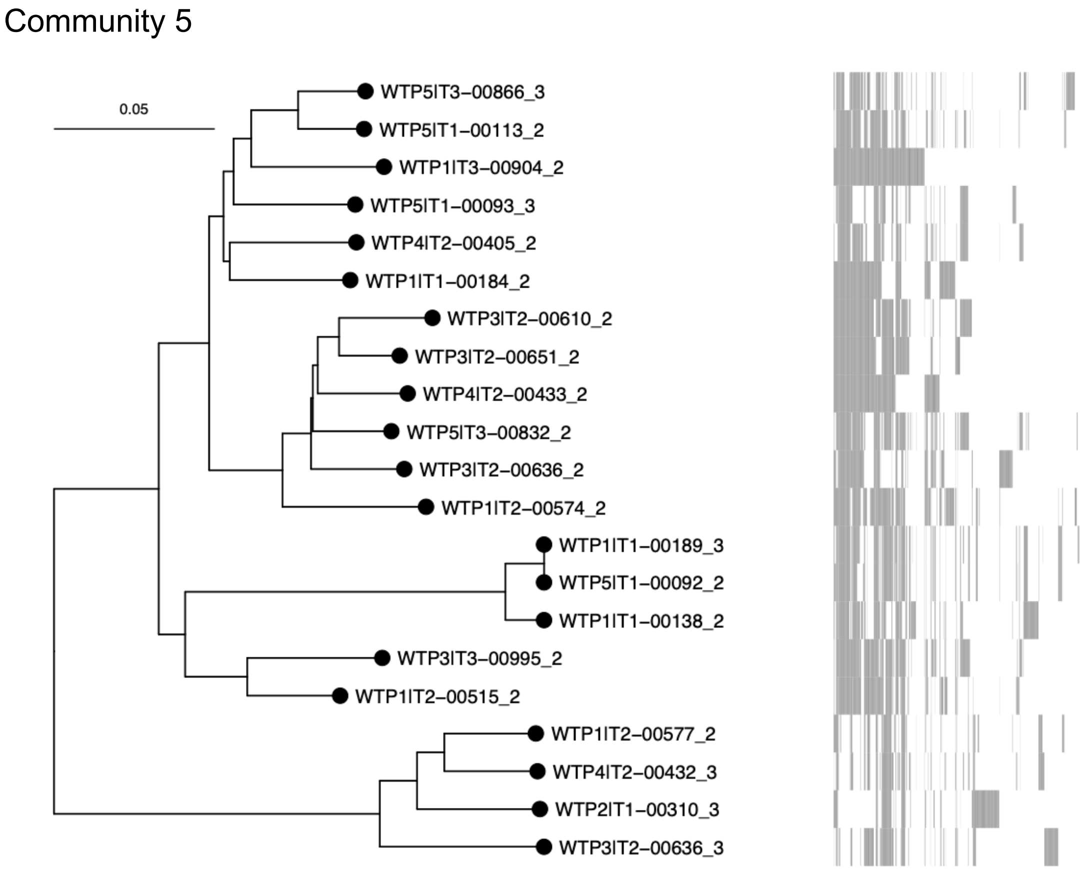


**Figure S6. Plasmid core-gene phylogeny for community 5.**

Community 6

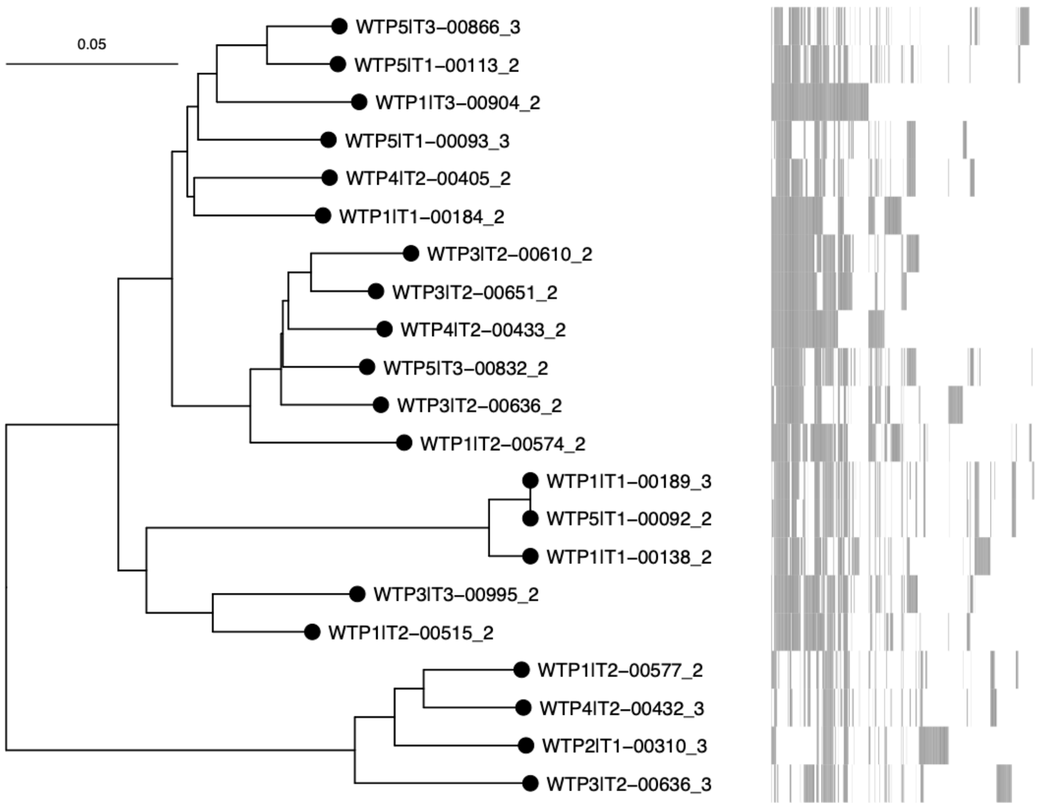


**Figure S7. Plasmid core-gene phylogeny for community 6.**

Community 7

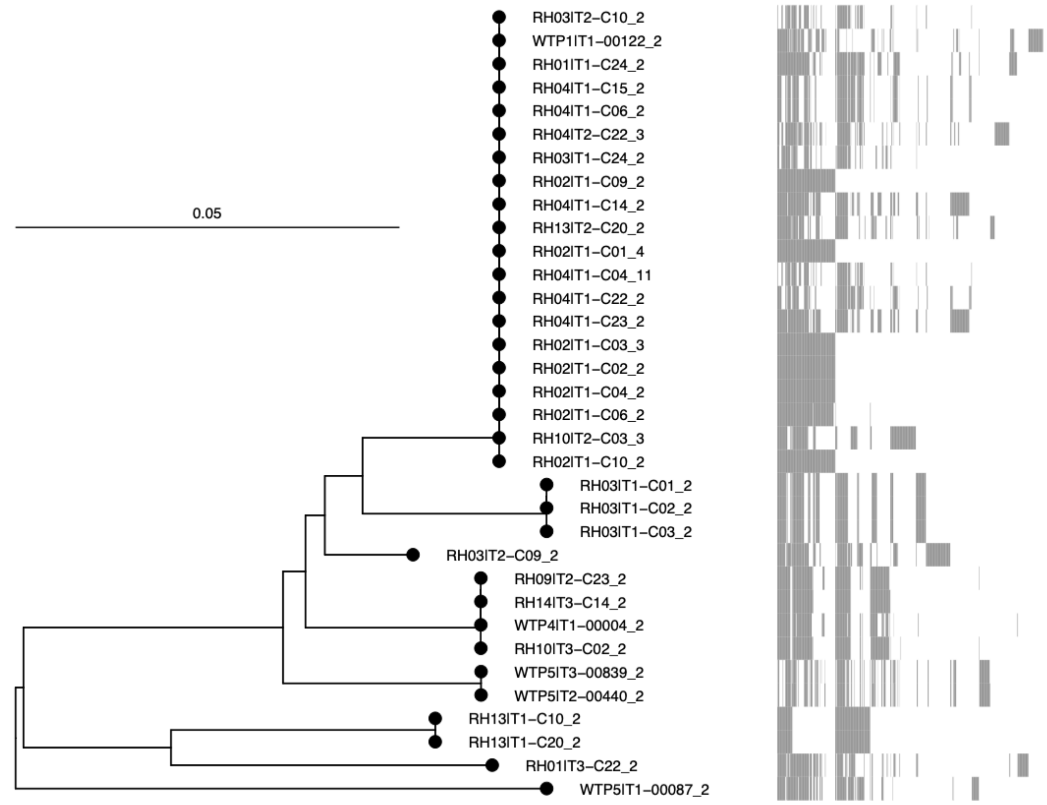


**Figure S8. Plasmid core-gene phylogeny for community 7.**

Community 8

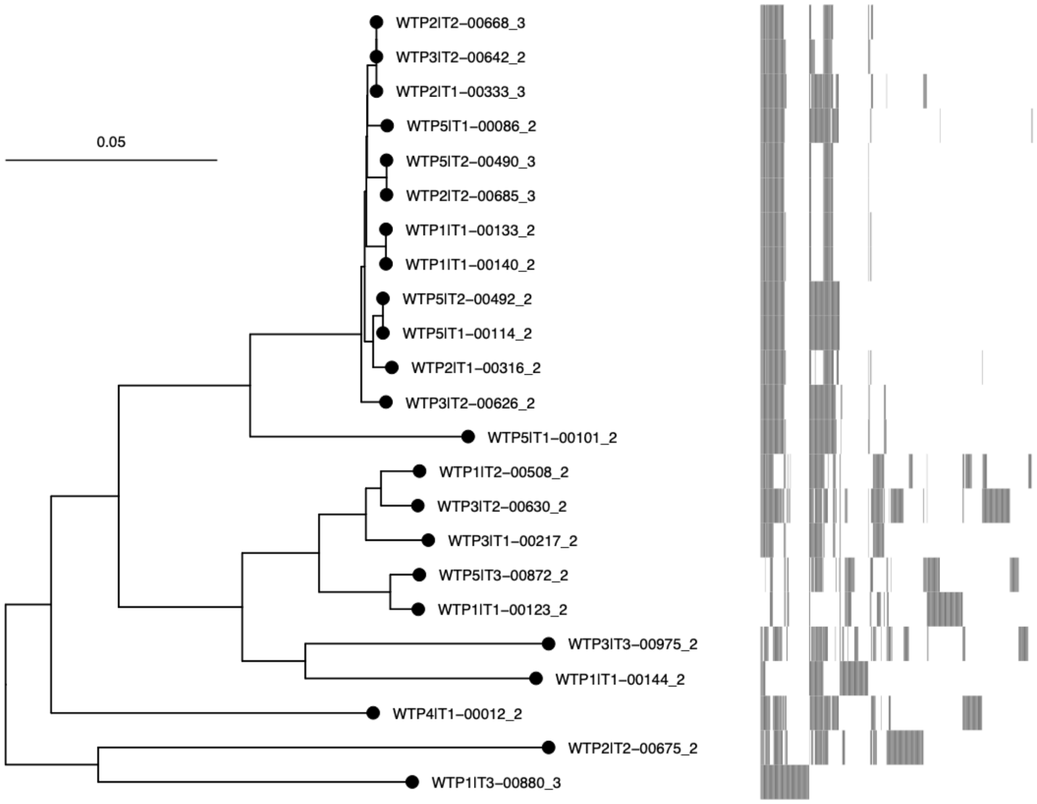


**Figure S9. Plasmid core-gene phylogeny for community 8.**

#### Community 9

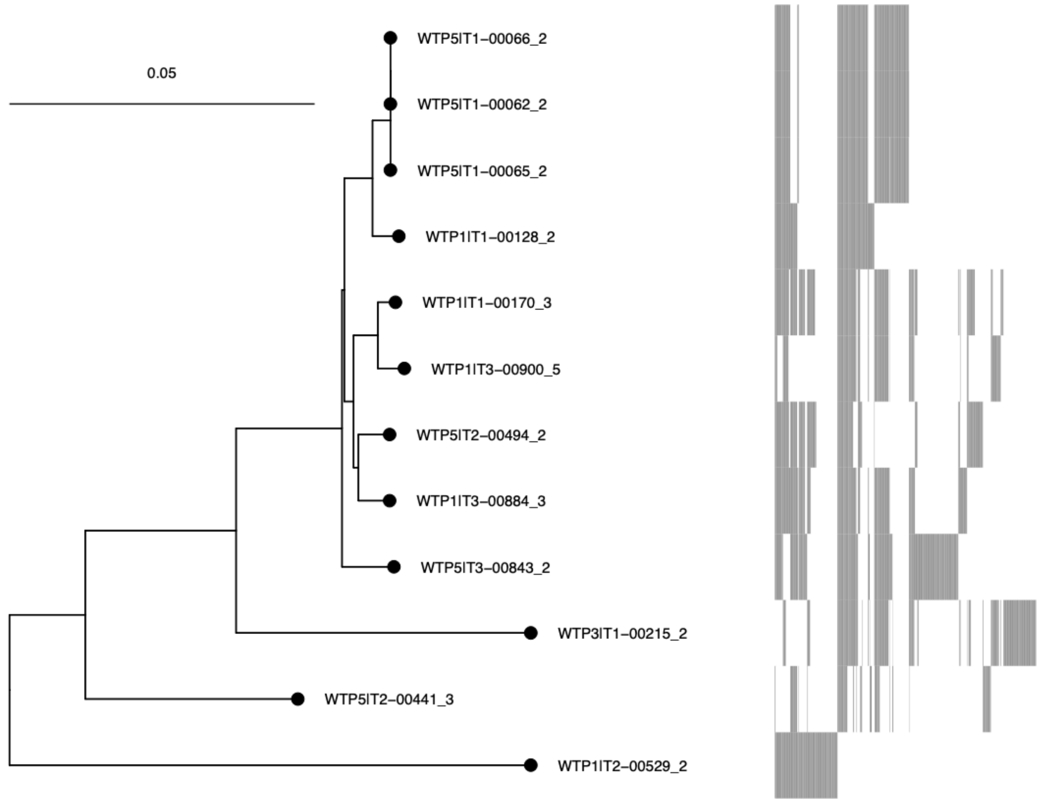


**Figure S10. Plasmid core-gene phylogeny for community 9.**

Community 11

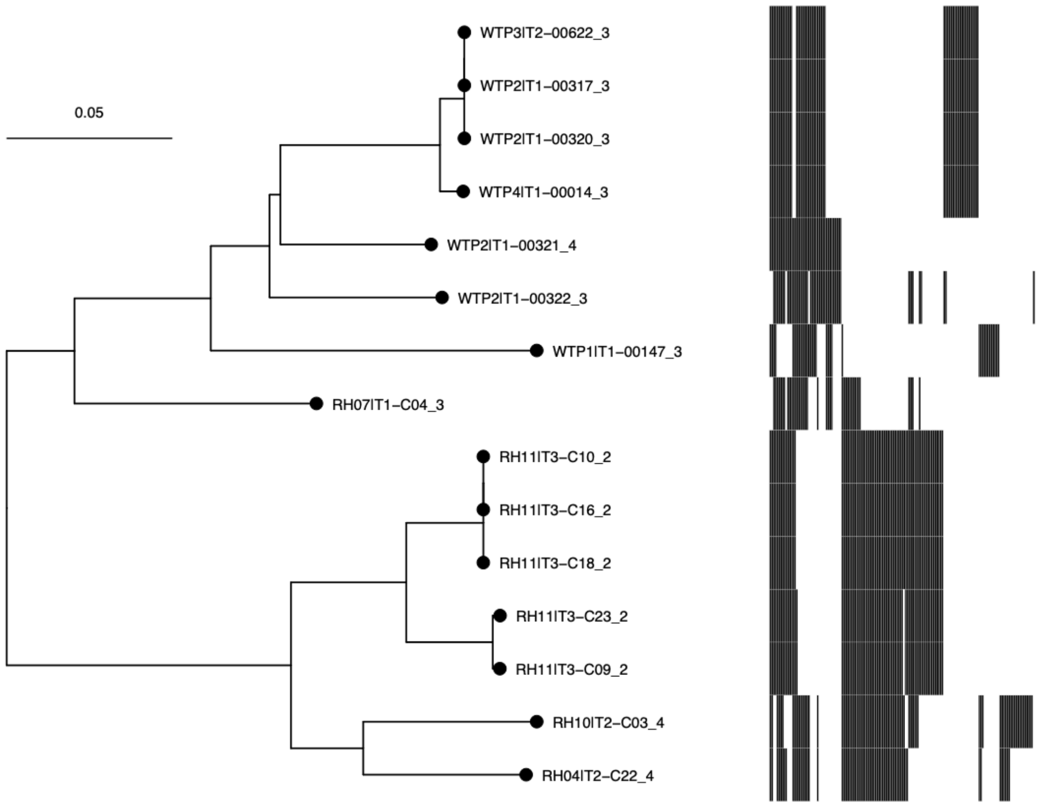


**Figure S11. Plasmid core-gene phylogeny for community 11.**

#### Community 13

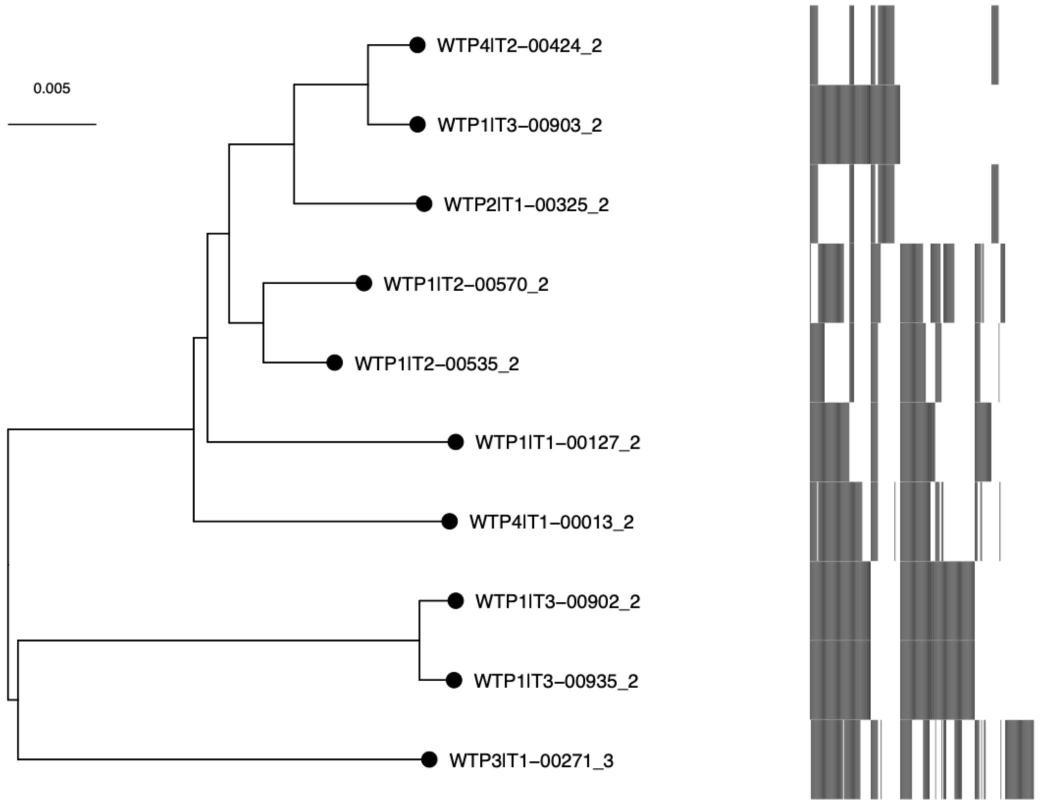


**Figure S12. Plasmid core-gene phylogeny for community 13.**

| Table S1: Replicon Haplotype Frequencies | | | |
| --- | --- | --- | --- |
| **Replicon Type(s)** | **Count** | **Replicon Type(s)** | **Count** |
| IncFIB, IncFII, IncFIIA | 156 | IncFII, IncFIIA, IncI1 | 2 |
| IncFII, IncFIIA | 104 | IncFIA, IncY | 2 |
| IncFIB, IncFII | 91 | IncFII, rep_cluster_48, rep_cluster_312, rep_cluster_959 | 2 |
| IncFIB | 65 | IncA/C2, IncFII, rep_cluster_959 | 1 |
| IncFIA, IncFIB, IncFII, IncFIIA | 58 | ColRNAI_rep_cluster_1987, IncFIA | 1 |
| IncFII | 30 | IncFIA, IncR | 1 |
| IncFIA, IncFIB | 26 | ColRNAI_rep_cluster_1291, IncFIA, IncFIB, IncFII | 1 |
| IncFII, rep_cluster_959 | 23 | IncFIB, IncH, rep_cluster_1150 | 1 |
| IncFIIA | 18 | IncFIB, IncP | 1 |
| IncFII, IncI1 | 16 | IncFIB, IncFII, IncP | 1 |
| IncFIA, IncFII, IncFIIA | 10 | IncFIB, IncR | 1 |
| IncFIA, IncQ1 | 9 | IncFIB, IncU, rep_cluster_1150 | 1 |
| IncFIA, IncFIB, rep_cluster_1760 | 9 | IncFIB, IncFII, IncFIIA, rep_cluster_1304 | 1 |
| IncFII, rep_cluster_1418 | 9 | IncFIB, IncFII, IncFIIA, IncH, rep_cluster_1304 | 1 |
| IncFIA, IncFII | 8 | IncFIA, IncFIB, IncFII, IncFIIA, rep_cluster_1704 | 1 |
| IncFIB, rep_cluster_1150 | 7 | IncFIB, rep_cluster_1804 | 1 |
| IncFIB, IncFII, IncI1 | 7 | IncFIB, IncFII, IncR | 1 |
| IncFII, IncR | 6 | IncFII, IncFIA, IncR | 1 |
| IncFIB, IncFII, IncFIIA, IncQ1 | 5 | IncFII, rep_cluster_312 | 1 |
| ColRNAI_rep_cluster_1291, IncFIB, IncFII, IncFIIA | 5 | IncFIB, IncFII, rep_cluster_48, rep_cluster_312 | 1 |
| IncFIA, IncFII | 4 | IncFII, rep_cluster_48, rep_cluster_959 | 1 |
| IncFIA, rep_cluster_1418 | 4 | IncFIIA, IncQ1 | 1 |
| IncFIA, IncFIB, IncFII, IncI1 | 3 | IncFII, IncFIIA, IncR | 1 |
| IncFIB, IncFII, rep_cluster_312 | 3 | IncFIA, IncFII, IncFIIA, IncX1 | 1 |
| IncFIA, IncFII, rep_cluster_959 | 3 | IncFII, IncFIIA, IncX1 | 1 |
| ColRNAI_rep_cluster_1987, IncFIB, IncFII | 3 | IncFIB, IncFII, IncFIIA, IncX1 | 1 |
| IncFIA | 2 | IncFIB, IncY | 1 |
| ColRNAI_rep_cluster_1291, IncFIA, IncFIB, IncFII, IncFIIA | 2 | ColRNAI_rep_cluster_1987, IncFII, rep_cluster_959 | 1 |
| IncFIB, IncFII, IncFIIA, IncR | 2 | IncFIA, rep_cluster_959 | 1 |
| IncFIB, IncFII, rep_cluster_48 | 2 | IncFII, IncN, rep_cluster_959 | 1 |
| IncFIB, IncFII, rep_cluster_959 | 2 | IncFII, rep_cluster_959, rep_cluster_1304 | 1 |

| Table S2: Permutation Test | | |
|---|---|---|
| Metadata Labels | Homogeneity *p*-value | Completeness *p*-value |
| Livestock, WwTP | *p*<0.0001 | *p*<0.0001 |
| Pig, Cattle, Sheep, WwTP | *p*<0.0001 | *p*<0.0001 |
| 14 Livestock Farms, WwTP | *p*<0.0001 | *p*<0.0001 |
| Livestock, 5 WwTPs | *p*<0.0001 | *p*<0.0001 |
| Livestock, Influent/Upstream, Effluent/Downstream | *p*<0.0001 | *p*<0.0001 |
| Host Genera | *p*<0.0001 | *p*<0.0001 |
| Time-point | 0.034 | 0.033 |

| Table S3: Community Core Gene Set Intersections | | | | | | | | | | | | | |
|---|---|---|---|---|---|---|---|---|---|---|---|---|---|
| | **1** | **2** | **3** | **4** | **5** | **6** | **7** | **8** | **9** | **10** | **11** | **12** | **13** |
| **1** | 3 | | | | | | | | | | | | |
| **2** | 0 | 4 | | | | | | | | | | | |
| **3** | 1 | 1 | 35 | | | | | | | | | | |
| **4** | 0 | 1 | 0 | 2 | | | | | | | | | |
| **5** | 0 | 0 | 0 | 0 | 2 | | | | | | | | |
| **6** | 1 | 0 | 0 | 0 | 0 | 13 | | | | | | | |
| **7** | 0 | 1 | 0 | 2 | 0 | 0 | 2 | | | | | | |
| **8** | 0 | 0 | 2 | 0 | 0 | 0 | 0 | 27 | | | | | |
| **9** | 0 | 0 | 0 | 0 | 0 | 0 | 0 | 13 | 18 | | | | |
| **10** | 0 | 0 | 0 | 0 | 0 | 0 | 0 | 0 | 0 | 0 | | | |
| **11** | 1 | 0 | 2 | 0 | 0 | 1 | 0 | 0 | 0 | 0 | 62 | | |
| **12** | 1 | 1 | 10 | 0 | 0 | 2 | 0 | 0 | 0 | 0 | 7 | 68 | |
| **13** | 1 | 1 | 7 | 1 | 2 | 2 | 1 | 1 | 0 | 0 | 21 | 15 | 88 |

| Table S4: Community Accessory Gene Set Intersections | | | | | | | | | | | | | |
|---|---|---|---|---|---|---|---|---|---|---|---|---|---|
| | 1 | 2 | 3 | 4 | 5 | 6 | 7 | 8 | 9 | 10 | 11 | 12 | 13 |
| 1 | 441 | | | | | | | | | | | | |
| 2 | 405 | 520 | | | | | | | | | | | |
| 3 | 375 | 395 | 463 | | | | | | | | | | |
| 4 | 347 | 344 | 343 | 419 | | | | | | | | | |
| 5 | 381 | 382 | 355 | 345 | 485 | | | | | | | | |
| 6 | 378 | 436 | 372 | 345 | 418 | 790 | | | | | | | |
| 7 | 395 | 455 | 409 | 366 | 393 | 509 | 638 | | | | | | |
| 8 | 368 | 399 | 366 | 334 | 385 | 429 | 405 | 505 | | | | | |
| 9 | 384 | 437 | 379 | 350 | 405 | 570 | 519 | 424 | 688 | | | | |
| 10 | 305 | 304 | 300 | 313 | 314 | 310 | 311 | 302 | 314 | 346 | | | |
| 11 | 131 | 131 | 132 | 142 | 134 | 132 | 142 | 131 | 138 | 128 | 151 | | |
| 12 | 272 | 282 | 263 | 263 | 255 | 255 | 263 | 250 | 254 | 246 | 123 | 316 | |
| 13 | 184 | 185 | 181 | 188 | 212 | 203 | 190 | 190 | 199 | 186 | 99 | 167 | 243 |

| Table S5 | | | | | |
|---|---|---|---|---|---|
| WwTW | Population equivalient (PE) | Primary treatment | Secondary treatmant | Tertiary treatment | Consented Flow ($m^3$/d) |
| WTP01 | 223,435 | PSTs | ASP | N/A | 50,985 |
| WTP02 | 49,522 | PSTs | ASP | Disc filters | 11,883 |
| WTP03 | 37,731 | PSTs | ASP | Sand filters | 11,476 |
| WTP04 | 26,905 | PSTs | Filters | N/A | 6,250 |
| WTP05 | 2,841 | PSTs | Filters | N/A | 2,000 |
